# Non-Equilibrium Spatial Encoding of Nanoscale Mechanical Relaxation in Growing Plant Epithelial Cells

**DOI:** 10.64898/2026.03.18.712596

**Authors:** Jacob Kienast, Ian Moore, Sonia Contera

## Abstract

A central problem in soft and biological physics is how molecular-scale activity and remodelling coarse-grain into emergent mechanical laws at larger scales. In *growing* cell walls (polymeric composite materials that surround 90% of living organisms’ cells) irreversible deformation is not controlled by elastic stress alone. Instead, growth depends on the interplay between energy storage, dissipation, and the local timing of viscoelastic relaxation. Although dynamic atomic force microscopy (AFM) resolves storage and loss moduli (*E′, E″)* of living walls at nanometre resolution, these observables have remained phenomenological and disconnected from constitutive field variables. Here we introduce a physics-based inversion framework that converts AFM measurements of epidermal cells of living Arabidopsis plants into spatially resolved fields of stiffness *k*, viscosity *η*, and relaxation time *τ*. By analysing the spatial gradients of E′ and E″, we uncover organized mechanical heterogeneities governed by cellular confinement and stress focusing. We demonstrate that the local relaxation time is encoded directly in the coupling between storage and dissipation, yielding the pointwise relation *τ* = (1/*ω*) ∂*E*’/∂*E*’’, where *ω* is the indentation frequency. This relation enables model-independent extraction of mechanical timescales and establishes a general route from nanoscale non-equilibrium rheology to continuum descriptions of growth in living and active soft materials.

**Significance:** How molecular-scale activity gives rise to tissue-scale form is a central challenge in biological physics. Although growth is fundamentally a non-equilibrium mechanical process, experimental measurements at the nanoscale have not been directly connected to the constitutive parameters that govern morphogenesis. We introduce a framework that converts dynamic atomic force microscopy maps of storage and loss moduli into spatially resolved fields of stiffness, viscosity, and relaxation time in living cell walls. By revealing that mechanical relaxation is encoded in the local coupling between elastic storage and viscous dissipation, our work provides a route from nanoscale rheology to growth-relevant mechanical timing. This establishes a quantitative bridge between molecular remodeling and continuum mechanics, enabling direct experimental constraints on multiscale theories of morphogenesis.

## 1. Introduction

Living systems generate hierarchical structure by converting molecular-scale activity into organized mechanical deformation across scales. Morphogenesis is thus a fundamentally non-equilibrium process in which energy consumption and material remodeling produce irreversible shape change. A central problem in soft and biological physics is how such microscopic activity coarse-grains into emergent constitutive laws that govern tissue-scale mechanics. In particular, polarized growth requires the coordinated regulation of stress, dissipation, and relaxation within mechanically heterogeneous materials. Understanding how these local viscoelastic processes encode large-scale form remains an open challenge. [1–5].

In plants, organ growth is strongly shaped by the epidermis, which behaves as a load-bearing “biomechanical shell” that integrates growth across scales [3, 5–7], Plant morphogenesis is therefore constrained (and enabled) by the primary cell wall: a living, fibre-reinforced composite whose macroscopic mechanics emerge from nanoscale interactions among cellulose microfibrils, matrix polysaccharides, and wall-associated proteins[8–11]. Genetics, biophysics, and modelling have converged on a mechanochemical-feedback picture in which wall composition and architecture pattern stresses, while stresses guide wall remodelling and anisotropy, coordinating growth from cell to tissue scales[2, 5, 12, 13] In this physics-to-molecules view, wall stress and biochemical loosening cooperate to set extensibility and growth anisotropy, dynamically tuned in vivo by cellulose synthesis, pH regulation, and hormone signalling[9, 14, 15]..]. Coen and Cosgrove further proposed that plant form emerges from feedbacks linking patterned gene activity, stress fields, and wall remodelling, emphasizing anisotropy and the need for constitutive laws valid across scales [3].

Ultimately, organ shape is set by the growth of individual cells. Growth is driven by largely isotropic turgor pressure, while directional expansion requires molecular structuring of the wall to maintain integrity [16–19]. Wall mechanics arise from coupled biochemical and biomechanical contributions that are actively modulated by enzymatic activity, hormonal regulation, and evolving structural anisotropy [8, 10, 11, 15, 20, 21] Consistent with this, changes in wall mechanics track morphogenesis [20, 22, 23] (reviewed in [24]), and tissue-level assays reveal strongly nonlinear behaviour that constrains models of microfibril sliding, reorientation, and extensibility [25, 26] In parallel, key wall-sensing pathways have been identified, including edge-localized RLP4/RLP4-L1 modules coordinating directional growth [27], and the SCOOP18–MIK2 ligand–receptor pair linking wall damage to immunity [12].

Despite this progress, a quantitative link between cell-wall mechanics and growth remains elusive. Available assays probe different physical quantities under distinct loading geometries and timescales—from tensile stretching and osmotic approaches to optical methods such as Brillouin microscopy and indentation-based AFM/CFM—and therefore often return non-equivalent “moduli”. [15, 24, 28]]. A recurring tension is that indentation-derived elastic moduli, frequently reported under diverse perturbations, are orders of magnitude lower than in-plane tensile stiffnesses that more directly constrain expansive growth, complicating their interpretation [15, 24] Moreover, most plant-wall AFM remains quasi-static and Hertz-fit-based, limiting access to subcellular time-dependent behaviour, although dynamic nanoindentation strategies have begun to address this gap [24, 29, 30]. Consequently, the field increasingly calls for integrating wall mechanics with cytoskeletal dynamics and trafficking in predictive multiscale models, and for cross-calibrating AFM, Brillouin, extensometry, and spectroscopic readouts to reconcile technique-dependent viscoelastic parameters [5, 13, 31, 32]

A further and crucial challenge is that growth is inherently dissipative: stress patterns alone are insufficient because walls both store and dissipate mechanical energy during deformation and remodelling [15, 16, 19]. Accurate descriptions of growth therefore require access to both elastic (stored) and viscous (dissipated) contributions and their associated time responses [29, 33] We previously showed that multifrequency contact-resonance AFM enables quantitative in vivo mapping of storage (E’) and loss moduli (E’’) in Arabidopsis cell walls, extracting viscoelastic parameters pixel-by-pixel directly from dynamic observables and providing an internal validity test of the assumed mechanical model [34] Together with related multifrequency/dynamic AFM developments [35–37], these approaches suggest a route to connect nanoscale rheology to growth-relevant wall physics and to functional phenotypes, as we showed with wall fucosylation [38] Because active wall remodelling and synthesis operate on longer timescales than high-frequency AFM probing, combining dynamic methods with controlled perturbations can help disentangle polymer-network mechanics from slower biochemical processes, as discussed in [34]

Here we extend our framework to resolve spatial fields of stress and dissipation across Arabidopsis epidermal cells spanning distinct geometries and organs. We find that dissipation landscapes do not simply track static stress: the gradients of stored (E’) and dissipated (E’’) mechanical energy systematically deviate from purely geometric predictions, revealing structured viscoelastic heterogeneity. From these measurements we derive a local inversion that extracts stiffness, viscosity, and relaxation time directly from AFM observables, and show that the mechanical time response scales with the gradient of E’ relative to E’’. Geometry-driven stress focusing and spatial confinement emerge as primary regulators of these constitutive fields. Together, the results identify a general physical mechanism by which heterogeneous living materials tune relaxation dynamics through the local balance of elastic storage and dissipation, thereby linking non-equilibrium rheology to morphogenetic control.

## 2. Results and discussion

### 2.1 Multifrequency AFM quantitative mapping of mechanical properties E’ and E’’

Using multifrequency AFM we have obtained time-dependent mechanical properties (i.e. complex modulus) of the cell walls of epidermal cells with different growth patterns. We have used a contact resonance imaging method whereby the cantilever is permanently in contact with the sample while it is simultaneously oscillated at one or more of its harmonics; the feedback is done on the deflection of the cantilever following the method developed by Cartagena et al. [35]. This is based on a previous method [37]. A schematic of the technique is given in Figs. S1, S2 of the SI. The method is applicable to any general Maxwell linear viscoelastic material, as we demonstrated previously [34]. Using this method, it is possible to calculate the elastic (or storage) modulus (E’) and loss modulus (E’’) for every pixel of the image, using the formulas:

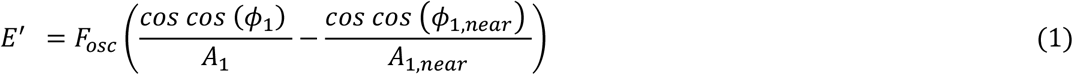

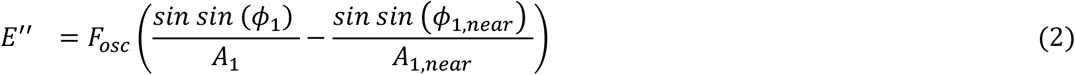

where *A*_1_ and *ϕ*_1_ are the amplitude and phase, respectively, of the cantilever, as extensively explained in [34]. The scaling factor, 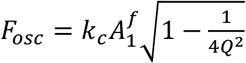 depends on the free amplitude 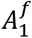 of the cantilever oscillation, the stiffness *k*_*c*_, and quality factor *Q* of the cantilever. Amplitude *A*_1,*near*_ and phase *ϕ*_1,*near*_, are measured at about ~ 10 nm above the surface to account for the hydrodynamic correction needed because the cantilever is oscillating in liquid [37, 39] (discussed in detail in [34]).

E’ quantifies the elastic mechanical energy stored in the sample (i.e., the elasticity) and is related to the displacement of the material, while E’’ measures the density of the energy dissipated during deformation (i.e. it is related to the viscosity). Fig. 1 shows maps of E’ and E’’ of at the junction of 4 epidermal cells of the hypocotyl, E’ and E’’ are over-imposed on the topography.

**Figure 1.**
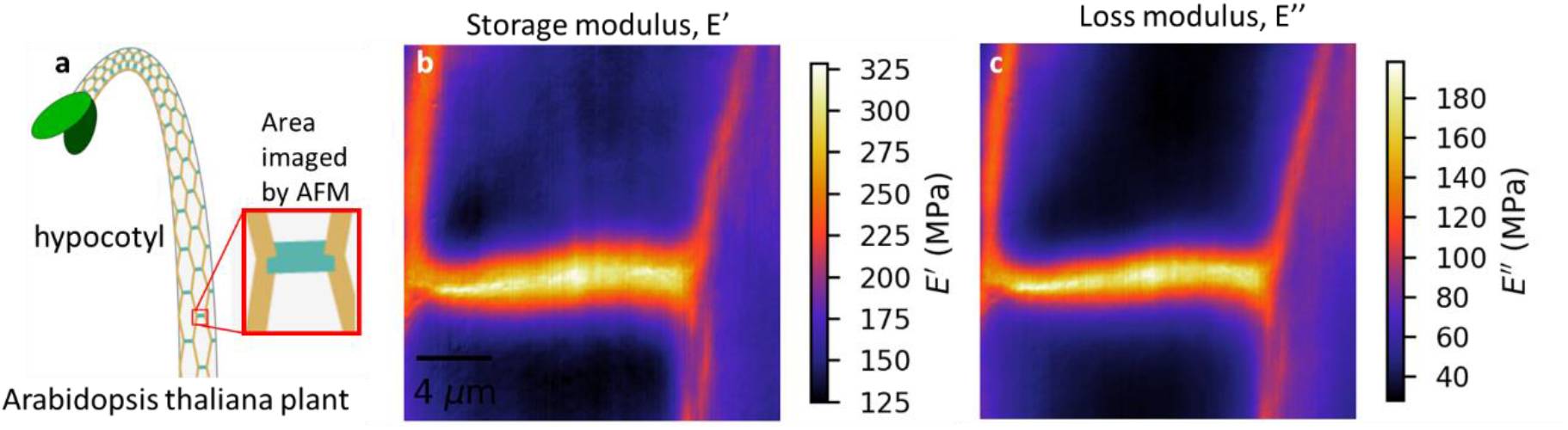
(a) shows a schematic of the plant hypocotyl and the area imaged by AFM. (b,c) show maps of elastic *E’* and loss *E’’* moduli of the scanned area calculated using equations (1) and (2) given in the text, respectively.

Fig.1 shows E’ and E’’ images of a junction of 4 cells at the surface of the hypocotyl of *Arabidopsis*, which agrees quantitatively with previously reported results using this technique and others as extensively discussed in [34].

In the SI Fig. S3, we demonstrate that the mechanical properties of the cell wall in our experiments match the standard linear solid viscoelastic behaviour, as expected from [34].

### 2.2 Nanoscale stress patterns and paths of energy dissipations in hypocotyl, and leaf (pavement and guard) cells in living Arabidopsis thaliana plants

Using the technique described in the *Materials section*, we used images of *E’* and *E’’* for different types of cells to calculate geometrical gradients (gradients of the z direction of the topography) 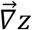, and of *E’* and *E’’* (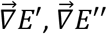); and compare them as they are over imposed on a topographical image of the cells. The geometrical gradient is also displayed in the image for comparison. This allows us to evaluate the relation between the geometry and the paths of stored (*E’*) and dissipated (*E’’*) energy of the cell walls. In Fig. 2, we show topographical images of epidermal cells with 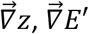, and 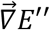 over-imposed on the topography for three types of cells.

**Figure 2.**
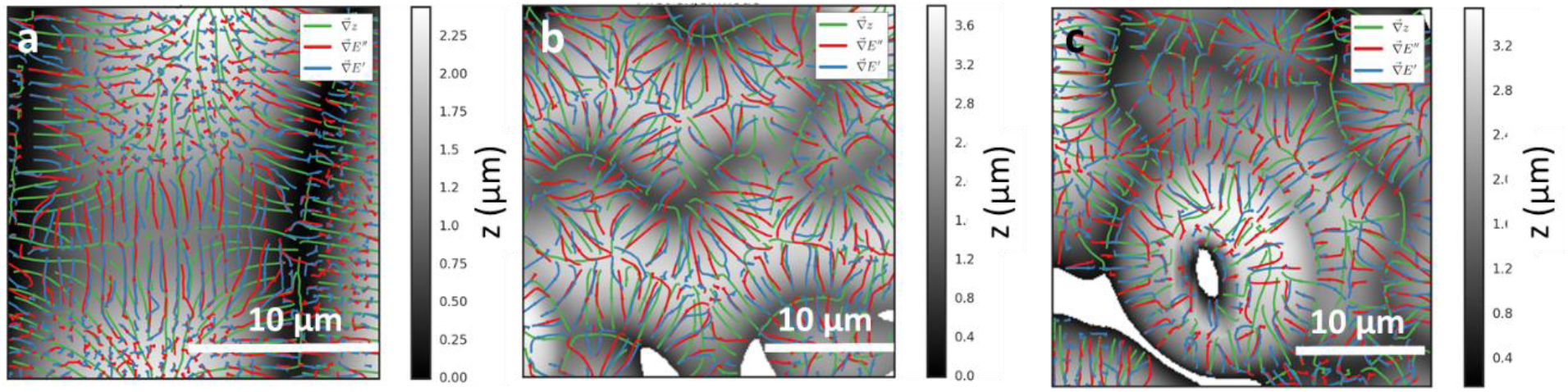
Gradients of geometry (green), storage modulus E′ (blue), and loss modulus E″ (red) overlaid on topographical images of Arabidopsis epidermal cells. (a) Four-cell junction (from Fig. 1), (b) leaf pavement cells, and (c) guard cells surrounding a stomatal pore. At regions of high geometric stress (cell junctions), E′ and E″ gradients align with the geometric gradient and are nearly parallel to each other; away from junctions, mechanical gradients decouple from geometry and from each other.

Fig. 2 (a), shows images of the gradients corresponding to the image of hypocotyl cells in Fig. 1, these cells are unidirectionally growing cells, as they occur in the embryonic part of the stem. The data clearly shows that the paths of stored 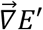 and dissipated energy 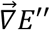 are strongly modulated by geometry in regions with high geometrical stresses, i.e., the cell junctions. At these locations, the mechanical gradients are either parallel or orthogonal to the spatial gradients, resembling the structures of geometrical stresses. However, away from these junctions, the paths of energy storage and dissipation are completely independent of the geometrical structures. Furthermore, it can be observed that paths of energy dissipation E’’ are not the same as for stored energy E’. This suggests a structurally different mechanism for these two phenomena and is in agreement with the current view of the plant cell wall and plant cell growth [9].

Figure 2 shows that discrepancies between the gradients of E′ and E″ and geometric stress patterns are largest at cell junctions, where mechanical stresses are expected to accumulate as immobile plant cells coordinate growth with their neighbours. This is reflected by elongated features at junctions, indicating rapid spatial variations in mechanical properties and elevated stress. Because these gradients do not align with the topographical gradient (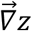), they do not arise from simple wall bending, but instead reveal stress accumulation specific to junctional regions. This observation refines earlier AFM studies reporting correspondence between elastic moduli and geometric stress [23, 40] by explicitly resolving dissipative contributions. We next examined two-dimensional growth in pavement cells, whose conserved lobed morphology underlies polarized organ expansion. Previous work showed that concave and convex regions experience distinct stress patterns and growth behaviours[2, 6, 23, 41, 42].

Figure 2b shows pavement cells in a leaf. As in the hypocotyl, the gradients 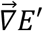 and 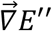 follow 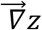 in regions of high geometric stress, particularly near cell junctions, but diverge from 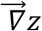 in regions of weaker geometric stress. Away from junctions, 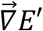 and 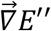 also follow distinct paths, indicating that elastic and dissipative contributions decouple when geometric constraints are reduced.

Pavement cell lobes arise from well-characterized mechanochemical processes, including ROP-mediated microtubule accumulation at future convex sites [43, 44], increased wall thickness, radial microfibril organization, and homogalacturonan demethylesterification, with a key role for self-expanding pectin nanofilaments[6] [45].. Turgor-induced buckling of anticlinal walls and localized reinforcement of periclinal walls have been proposed as drivers of lobe formation [41], while recent work suggests that specialized cortical microtubules spanning periclinal and anticlinal walls pattern anisotropic cellulose domains at sites of elevated tensile stress, nucleating lobes mechanically coupled through the pectin-rich middle lamella[2].

Our measurements show that in regions of high geometric stress, elastic (E′) and dissipative (E″) gradients coincide, indicating a local balance between stored elastic energy and mechanical dissipation. This supports current models linking lobe formation to the interplay between elastic components such as cellulose and more dissipative pectin-rich matrices. At the same time, it raises the question of how the balance between E′ and E″ shapes cell geometry. Addressing this requires access to the local time response, which ultimately governs how material properties couple to shape and is examined in Section 2.3.

Finally, we examined guard cells, the paired epidermal cells that regulate stomatal pore opening and gas exchange (Fig. 3c). Guard cell walls are thought to be mechanically specialized to withstand high turgor while enabling reversible deformation, and recent work has established their central role in stomatal development and dynamics [46]. Because stomatal opening and closing are reversible, guard cells are expected to behave predominantly elastically.

**Figure 3.**
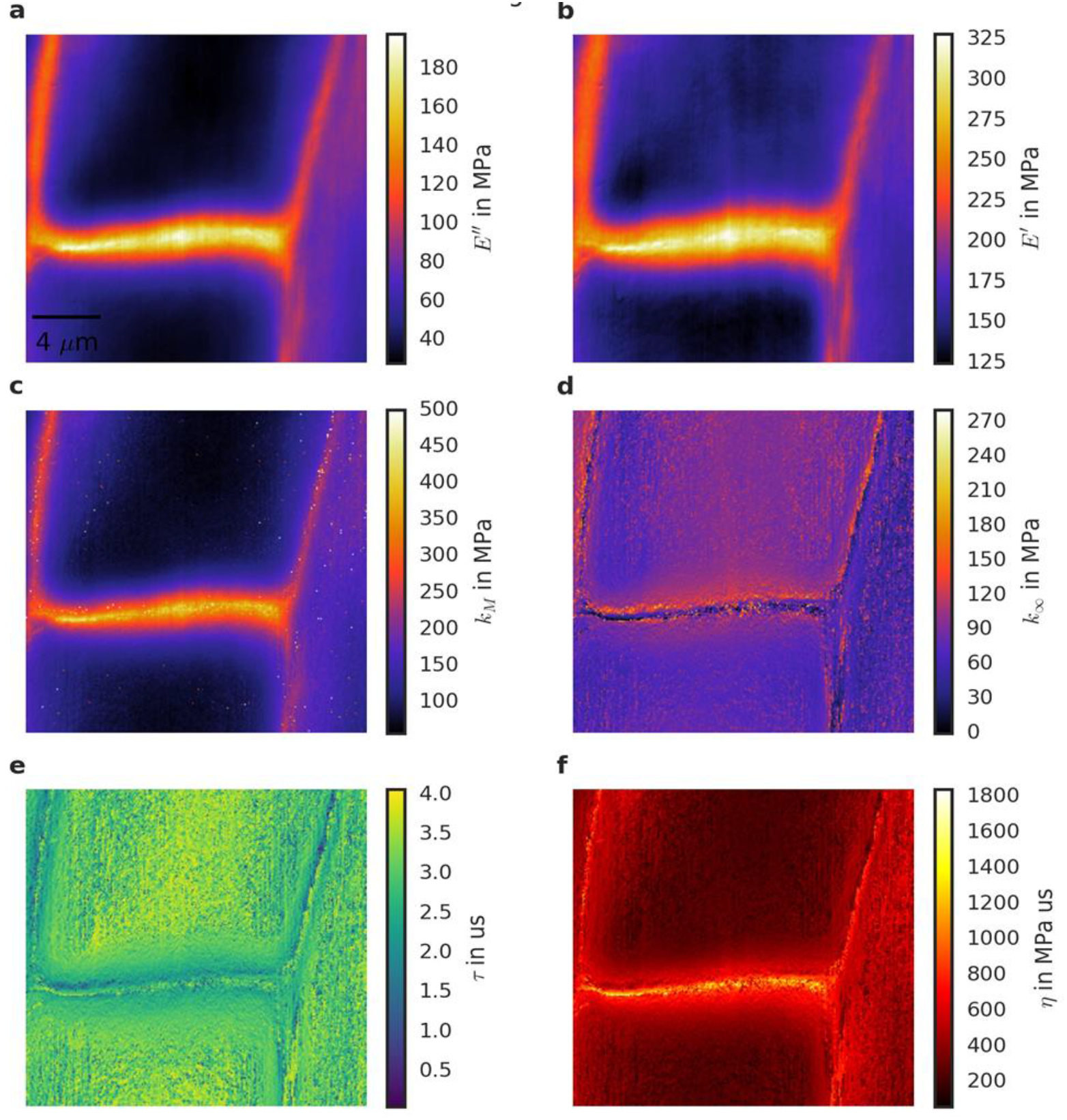
Standard linear solid viscoelastic quantities in hypocotyl cells. (a,b) show *E’’* and *E’* for the scan, which is already shown in Figs. 1, 2; (c-f) show *k*_*M*_, *k*_*∞*,_ τ, and η, respectively.

Consistent with this expectation, mechanical gradients align with geometric stress gradients at regions of high stress, particularly near cell junctions (Fig. 3c). Away from these junctions, however, gradients of stored (E′) and dissipated (E″) energy decouple from geometry and from each other, indicating distinct elastic and dissipative pathways. This behaviour mirrors recent AFM and modelling studies reporting polar stiffening of stomatal complexes in Arabidopsis and other species [47], which we previously confirmed using dynamic AFM [38]. Carter *et al*. [47], proposed that polar stiffening arises from pectin-mediated mechanical pinning at the guard cell ends, enhancing stomatal sensitivity to turgor pressure and challenging models based solely on radial wall thickening.

Our results show that both E′ and E″ gradients are enhanced at the poles, raising the question of how their balance determines the local time response and whether elastic or viscous contributions dominate. We address this directly in the next section.

### 2.3 Nanoscale maps of viscosity, stiffness, and local time response in hypocotyl, pavement and guard cells

To compute the local time response (relaxation time) τ of the cell wall, we require a constitutive model linking the storage (E′) and loss (E″) moduli to τ. Linear viscoelastic models—including the Kelvin–Voigt, Maxwell, and Standard Linear Solid (SLS)—represent materials as combinations of Hookean springs and dashpots and allow extraction of viscoelastic parameters such as stiffness k, viscosity η, and relaxation time τ at a given frequency. We previously showed that the mechanical response probed here is well described by the SLS model (see [34] and SI section S1), enabling pixel-resolved determination of τ and the associated viscoelastic parameters from AFM images.

This would be straight forward if the underlying model was the Maxwell model (one spring and one dashpot in series, which describes fluid behaviour), where the *relaxation time τ* is given by the ratio of the elastic and the loss modulus *ωτ = E’/E’’*. In the case of SLS this is more complicated, since in this case, *E*^′^ = *k*_∞_ + *E*^′′^*ωτ* (see section S2 in the SI)

From this equation it can be readily shown that *τ* for each pixel can be obtained using the relation:

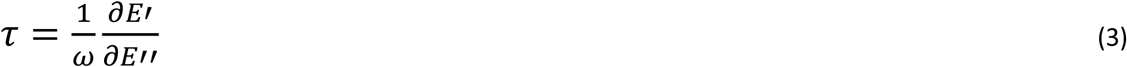

Namely, the time response locally is related to the gradients of the elastic energy stored E’ vs the energy dissipated E” at a given frequency. Interestingly, a similar mathematical approach for a seemingly unrelated problem was used by Miyahara, et al [22], who calculated a quantum mechanical tunnelling rate for the atomic orbitals of quantum dots using conservative and dissipative quantities.

This gives us a simple method to map local time responses. In the SI section S2 we show a numerical method to calculate maps easily from the experimental data using equation (3) and the AFM observables.

Using our approach, it is now possible also to calculate the rest of the viscoelastic quantities of the standard linear solid, namely *k*_∞_, *k*_m_, and *η*, using the following equations

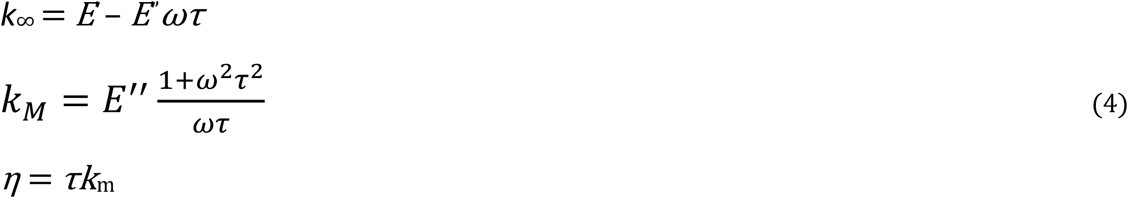

Using these equations and the numerical implementation described in SI section S2, we computed maps of the viscoelastic properties of Arabidopsis hypocotyl cells (Fig. 3). Anticlinal walls appear with sharper contrast than in the E′ and E″ maps because E′ and E″ represent energy densities that include internal wall stresses, which must vary continuously across wall interfaces. In contrast, the extracted viscoelastic parameters are not constrained by stress continuity, leading to sharper spatial transitions. With a pixel size of 78 nm and an estimated anticlinal wall thickness of 100–200 nm [23]; the images in *k*_m_, *τ*, and *η* in Fig. 3 seem to accurately reproduce the actual positions of those walls.

In this representation, not only cell walls but additional structures become apparent, particularly in Fig. 3 (e,f). These features are unlikely to reflect the molecular architecture of the wall, given the 78 nm pixel size and the much smaller dimensions of cellulose microfibrils (~5 nm) and matrix components. Instead, they likely represent a projection of effective mechanical properties over the penetration depth of the indentation and the oscillatory mechanical wave launched by the cantilever. While the indentation depth is known from the quasi-static indentation and it is about ~ 300nm (SI, Figure S1), the penetration depth *λ*_*p*_ can be estimated by the oscillation velocity ⟨*v*_0_⟩ = 1/2*A*_1_*ω* _1_and the relaxation time *τ*_*1*_, which gives *λ*_*p*_ = ⟨*v*_0_⟩*τ*_1_≈ *1*.*3nm*. The small value caused by the oscillation compared to the initial indentation makes this contribution negligible, and the mechanical properties measured in a pixel result mainly from the area that is in contact with the probe.

Finally, we can compare the derived *τ* and *k*_m_ with the fitted values from our previous publication [34] where these quantities were obtained from a different mathematical procedure by fitting the average of whole images. From Fig. 3, we obtain a mean (± SD) value of 2.9 ±0.5 µs, which is very close to the values obtained from the fit in our previous work, which where *τ*_*fit*_ = 2.36 µs to 2.57 µs. For *k*_m_ we obtain 144 ±73 MPa for the mean value for the local map and *k*_m_ = 110MPa to 226MPa. With these results we can conclude that the average behaviour of the spatial maps is in agreement with the average behaviour obtained by the fits of the data to the model.

A particularly relevant system to investigate stress dependency of elastic and viscous behaviour are the pavement cells in leaves because they have been shown to display a very specific stress pattern and that regions of concave (lobes) or convex (necks) curvature differ in growth [14, 33]. Below we use our technique to investigate pavement cells in the cotyledons. Figure 4 shows the topography of a typical structure of pavement cells from a cotyledon of a 4-day old Arabidopsis seedling.

**Figure 4.**
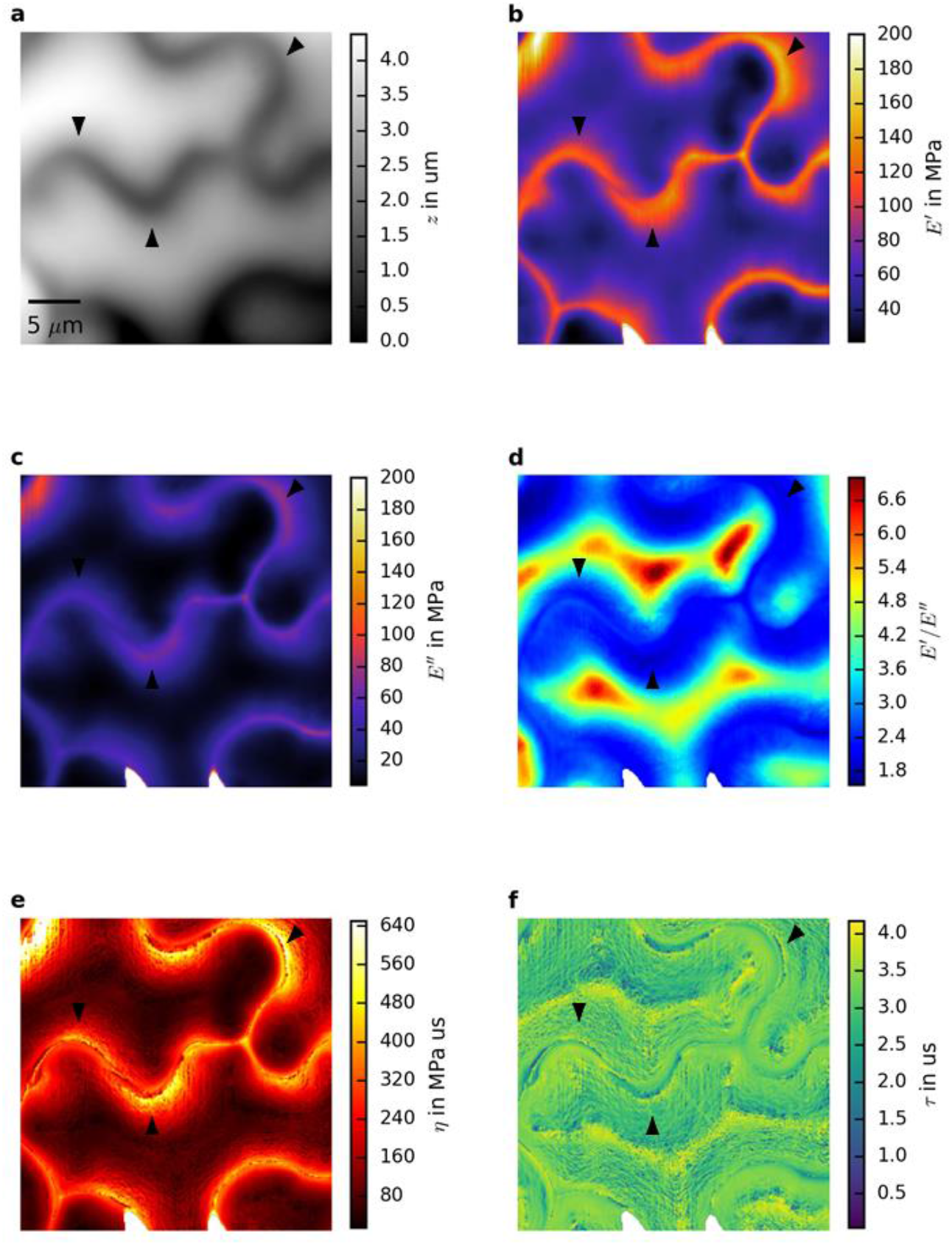
Standard linear solid viscoelastic quantities in pavement cells of Arabidopsis leaves. (a) shows the topography of the cells using AFM, (b) shows E’, (c) shows the corresponding E’’ map and (d) shows their ratio E’/E’’; (e-f) show η and τ, respectively.

Pavement cells exhibit a characteristic jigsaw-like geometry that generates localized stress concentrations at regions of concave curvature (arrowheads in Fig. 4). These geometric stress patterns are reflected in the E′ and E″ maps (Fig. 4b,c), confirming that the dynamic measurements are sensitive to both geometry-induced stress and intrinsic material properties. Compared with hypocotyl cells, pavement cells show substantially lower moduli (E′ ≈ 70 ± 23 MPa; E″ ≈ 26 ± 20 MPa), nearly an order of magnitude smaller, except at highly stressed junctions where values approach those of the hypocotyl. This difference likely reflects higher stress and mechanical reinforcement in rapidly elongating hypocotyl cells and/or softer, more isotropic wall architectures in pavement cells, where less aligned cellulose microfibrils may reduce effective stiffness.

Pavement cells also show a modestly higher E′/E″ ratio than hypocotyl cells (Fig. 4d; hypocotyl: ≈2.5 ± 1.0; pavement: ≈3.2 ± 1.0, mean ± SD), consistent with developmental differences in viscoelastic behaviour, as more mature cells tend to be less viscous and more elastic. Unlike the individual moduli, E′/E″ is similar at concave and convex cell regions. Viscosity maps (Fig. 4e) reveal lower average values in pavement cells (176 ± 80 MPa) than in the hypocotyl (408 ± 180 MPa), although this does not imply greater extensibility, which depends strongly on cell geometry [5, 48]. Spatial variations in viscosity are consistent with growth inhibition at necks and promotion at lobes, as previously proposed [6, 23, 43].

To interpret these results, we draw on theoretical models identifying viscosity as a key growth parameter tunable by the plant [16, 19, 33] In these models, viscosity is dominated by stress-dependent cross-link breakage, with the breakage rate *κ_off* increasing exponentially with cross-link stretching, and thus being highly sensitive to tissue stress and geometry.

In this framework, our data suggest that cross-links at lobes are under higher tension, while those at necks are more relaxed. Assuming comparable cross-link lengths, this interpretation is consistent with the higher cellulose microfibril density at necks inferred from microtubule patterning [14 Consequently, concave regions exhibit increased viscosity and reduced extensibility, whereas convex regions show reduced viscosity and enhanced extensibility. Here, viscosity reflects resistance to plastic deformation arising from cross-link dynamics.

While local variations in viscosity can be explained by cross-link breakage, the intrinsic viscosity of the polymer network requires separate consideration. From Fig. 4, we obtain an average relaxation time τ_1_ ≈ 3.1 ± 0.5 µs for pavement cells, comparable to that of the hypocotyl (2.9 ± 0.5 µs), consistent with a polymer-network response characterized by a broad, nearly continuous distribution of relaxation times.

Finally, we examined guard cells, which exhibit pronounced mechanical anisotropy during stomatal opening and closing as they transition between flaccid and turgid states. During inflation, guard cells must displace surrounding pavement cells, which remain turgid, suggesting anisotropic mechanics between the pore-facing and pavement-facing sides. Mechanical maps of a cotyledon pore (Fig. 5a) are consistent with this expectation and agree with our previous work showing that guard-cell time responses can be preserved in mutants that compensate reduced elasticity with increased dissipation [38].

**Figure 5.**
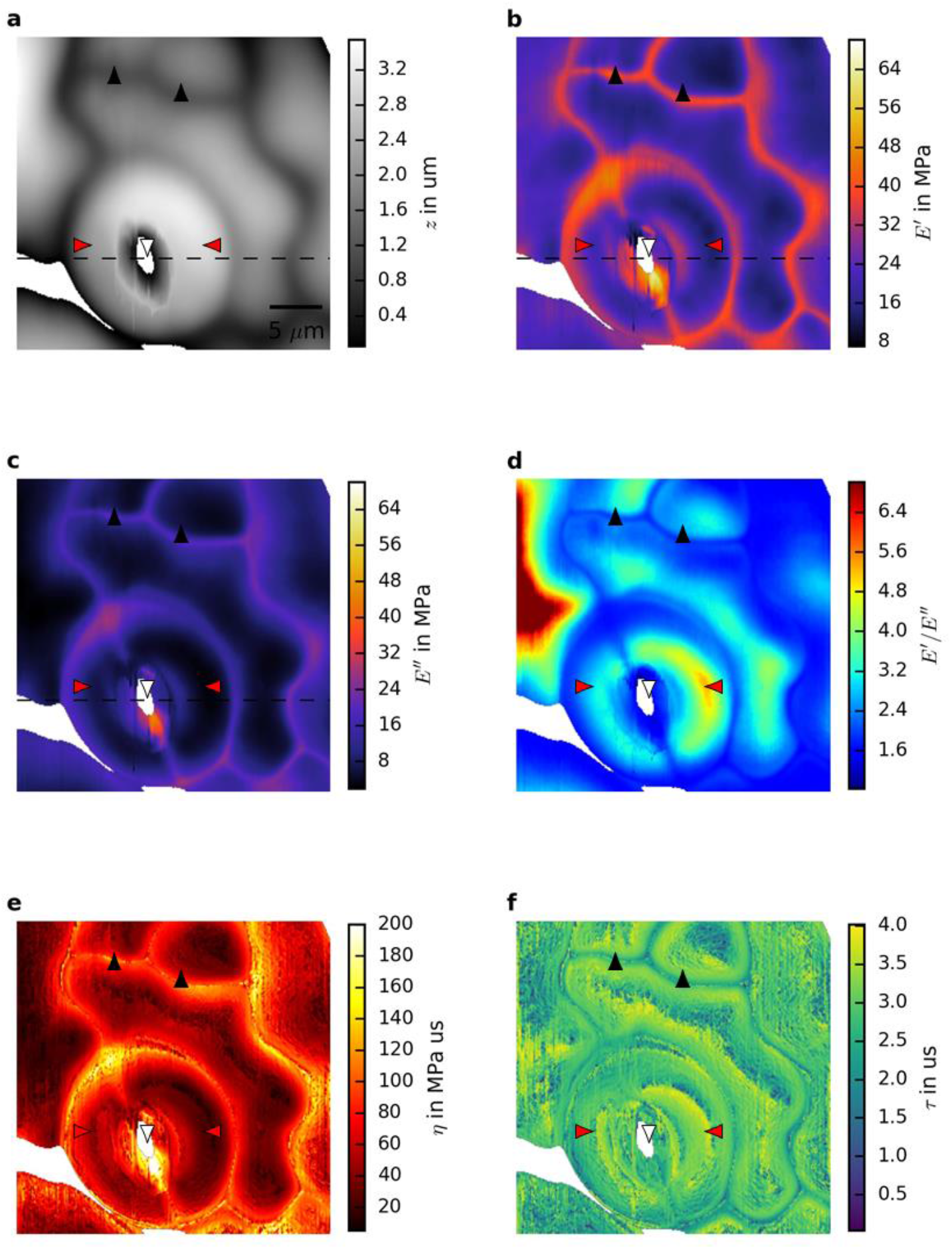
Viscoelastic properties of turgid guard cells with an open pore. (a) Topography of a stomatal pore embedded in pavement cells (white arrowhead, pore; red arrowheads, guard cells); dashed line marks the profile in Fig. 6. Storage (b) and loss (c) moduli reveal mechanical asymmetry between inner and outer guard-cell walls. (d) Inverse loss tangent is elevated at the apex. (e,f) Maps of viscosity and relaxation time.

To probe developmental sensitivity, we analysed a 5-day-old seedling, in which guard cells and surrounding pavement cells are not yet fully mature. The small pore diameter (~1.5 µm, compared with up to ~30 µm in mature stomata) and characteristic precursor cell shapes indicate an early developmental stage. Correspondingly, E′ and E″ values for pavement cells (Fig. 5b,c) are lower than in mature tissue (Fig. 4), with average E′ ≈ 25 ± 8 MPa and E″ ≈ 11 ± 7 MPa, consistent with previous quasi-static and dynamic indentation measurements [14, 29].

Radial profiles of topography, E′, and E″ (Fig. S4, SI) reveal a clear mechanical asymmetry across the guard cells, consistent with earlier reports of asymmetric guard-cell structure in open stomata [47].

## 3. Conclusions

In this work we establish a route from dynamic AFM observables to constitutive state variables that are typically inaccessible in nanoscale measurements. By combining spatial maps of the storage and loss moduli (E′, E″) with their gradients, we derive an experimentally implementable inversion that yields local stiffness, viscosity, and relaxation time (k_i_, η_i_, τ_i_) directly from imaging data. This promotes dynamic AFM from a phenomenological probe of moduli to a quantitative tool for extracting spatially resolved constitutive fields governing non-equilibrium deformation.

Although contact-resonance AFM operates at kHz frequencies, these measurements isolate short-time polymer-network modes that define the intrinsic relaxation spectrum of the wall. In hierarchical polymer composites, long-time yielding depends on these fast relaxation processes [34]. Consistently, we find that spatial variations in τ and η extracted at sub-millisecond timescales systematically track growth-relevant heterogeneity and confinement.

Across epidermal geometries, dissipation emerges as an organized component of the mechanical state rather than a passive by-product of stress. Spatial confinement strongly reshapes viscoelastic parameters, while active remodelling locally rebalances elastic storage and dissipation. Together, these results identify a general physical mechanism by which heterogeneous living composites tune relaxation dynamics to regulate irreversible growth.

More broadly, the framework provides a quantitative bridge between nanoscale rheology and continuum morphogenesis. By rendering dissipation measurable as a field variable, it supplies experimentally grounded parameters for multiscale models and advances toward predictive constitutive descriptions of growing active materials.

## 4. Materials and Methods

### Plant material and growth conditions

Arabidopsis thaliana ecotype Columbia (Col-0) wild-type plants were used for all AFM experiments. Seeds were surface-sterilized with 70% ethanol and grown vertically on Murashige and Skoog medium (2.2 g L^−1^; Sigma-Aldrich, pH 5.7) supplemented with 1% sucrose and 0.8% Bacto agar (BD Biosciences).

### AFM sample preparation

Seedlings were mounted on 15 mm metal probe holders using a thin layer of medical adhesive spray (Hollister 7730). Seedlings were positioned under the apical hook, incubated in a humid chamber for 15 min to allow adhesive setting, and subsequently covered with 50–100 µL of water for AFM imaging.

### AFM experiments and analysis

AFM measurements were performed using a Cypher ES microscope (Oxford Instruments Asylum Research) operated in contact-resonance mode with deflection feedback and photothermal actuation, as described previously [34] and summarized in SI Fig. S1. Nanosensors PPP-NCLAuD cantilevers (nominal spring constant ≈36 N m^−1^; resonance frequency in water ≈78 kHz; Q ≈ 9.7) were calibrated using the Sader method [49] Scans were acquired at 2.44 lines s^−1^ with 255 pixels per line (pixel size 78 nm). The free oscillation amplitude was set to ~14 nm, with an in-contact amplitude of ~4 nm and a deflection setpoint of 0.3 V, corresponding to an indentation depth of ~300 nm. The drive frequency was retuned before each scan, and the free-phase was set to 90°. Quasi-static indentation curves were collected at the end of each scan for calibration of amplitude and phase. A schematic of the experimental workflow and theoretical framework is provided in SI Fig. S2.

For guard cell measurements (Fig. 5), cotyledons from 5-day-old seedlings were mounted abaxial side up using the same adhesive protocol. Cotyledons were used due to the uneven surface of true leaves. Samples were maintained hydrated during adhesive setting, and half-strength MS medium was used during imaging. AFM data were analysed in Python 3.5 using previously described routines [34] (code available at https://github.com/jcbs/ForceMetric). Statistical comparisons were performed using two-sample t-tests.

## Acknowledgements

J.K., I.M., and S.C. acknowledge funding from the Biotechnology and Biological Sciences Research Council (BBSRC) under grant BB/P01979X/1.

## Data Availability

The data that support the findings of this study are available from the corresponding author upon reasonable request. The custom analysis code developed for this work is openly available at https://github.com/jcbs/ForceMetric.

