## Supplementary material for "Non-Equilibrium Spatial Encoding of Nanoscale Mechanical Relaxation in Growing Plant Epithelial Cells": Suplementary Material

**Contents:**

**Figures:**

- Fig. S1 Schematic representation of the multifrequency atomic force microscopy (AFM) technique used in this study, based on (Seifert et. al., 2021).
- Fig. S2: Processing flowchart including the experimental setup and calibration, and summary of assumptions necessary for calculating E’ and E’’
- Fig. S3: Validity of the standard linear solid test
- Fig. S4. Mechanical asymmetry in guard cells.

**Sections:**

- S1 Validity of the standard linear solid test
- S2 Theory to calculate the time response and numerical implementation $\tau$
- References

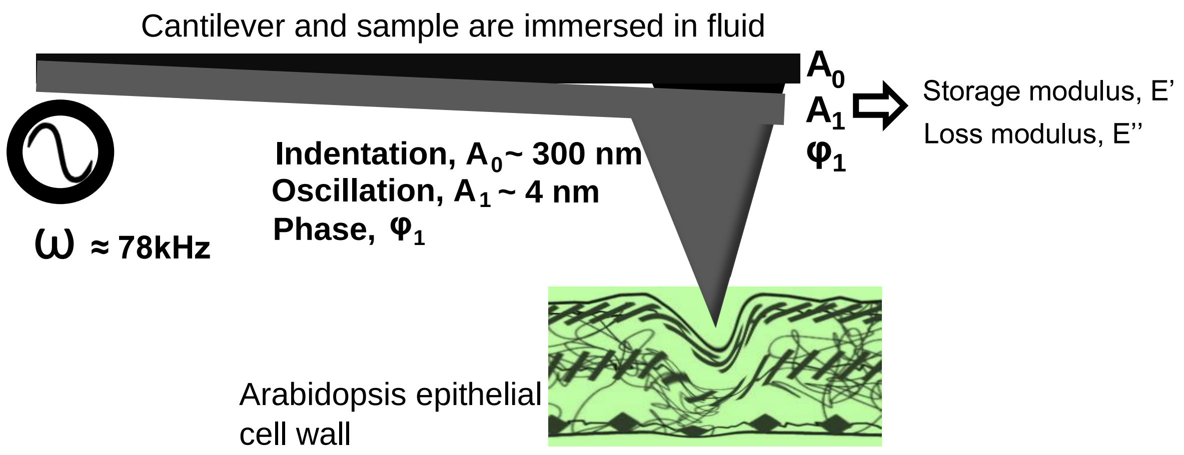


**Fig S.1 Schematic representation of the multifrequency atomic force microscopy (AFM) technique used in this study, based on ref (1). It is a contact resonance technique, which is done by producing a constant indentation of the cell wall of the leaves of living plants in medium, and a simultaneous oscillation of the cantilever. The imaging feedback is done on the deflection of the cantilever. Using a mathematical model independent of the tip geometry it is possible to produce nm-resolution maps of the elastic modulus *E'* and loss modulus *E''*. This technique has been shown to produce quantitatively correct values of time dependent mechanical properties of plant cell walls, unlike other indentation-based techniques where the cantilever is not in constant contact with the sample as we have extensively demonstrated in (1) The technique is summarised in the flowchart in Fig. S2.**

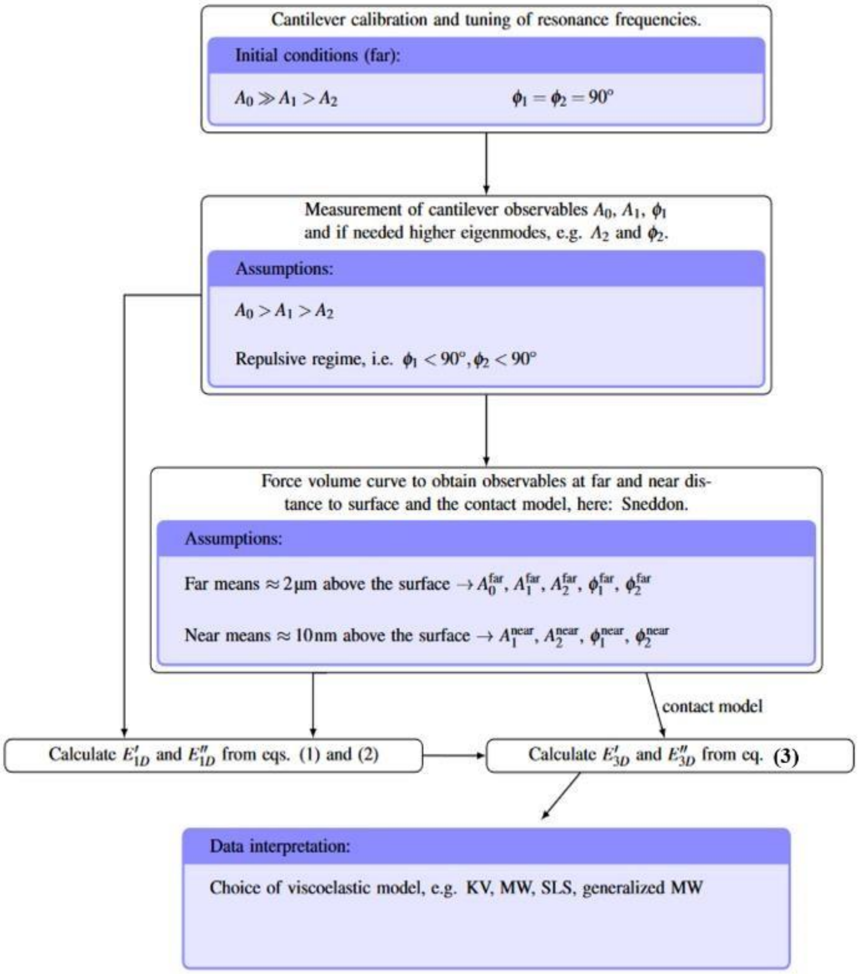


**Fig. S2: Processing flowchart including the experimental setup and calibration, and summary of assumptions necessary for calculating E' and E'' using the procedure explained in detail in the SI of reference (1) In the figure KV, stands for the Kelvin-Voigt model, MW is the Maxwell model, SLS is the standard linear solid. After calibration of the cantilever, the cantilever observables are measured. Then to be able to use the theory described in (1) we obtain the cantilever observables far and near the sample as outlined in the flowchart. Then we are able to calculate the 1-dimensional (1D) *E'* and *E''* using the equations below:**

$$F_0 = k_c A_0 = E_0(t)\delta_0$$

$$E' = F_{\mathrm{osc}}\left(\frac{\cos(\phi_1)}{A_1} - \frac{\cos(\phi_{1,\mathrm{near}})}{A_{1,\mathrm{near}}}\right)$$

$$E'' = F_{\mathrm{osc}}\left(\frac{\sin(\phi_1)}{A_1} - \frac{\sin(\phi_{1,\mathrm{near}})}{A_{1,\mathrm{near}}}\right).$$

**Then using the contact model (in this case Sneddon; see again the SI of ref. (1) for how this is determined we obtain *$E_0$* and then we can calculate the 3-dimensional *E'* and *E''* (3D) using the following equations:**

$$E'_{3D} = \frac{E'_{1D}}{2}\sqrt{\frac{1-\nu^2}{\tan(\alpha)}\frac{\pi}{2}\frac{E_0}{F_0}}$$
$$E''_{3D} = \frac{E''_{1D}}{2}\sqrt{\frac{1-\nu^2}{\tan(\alpha)}\frac{\pi}{2}\frac{E_0}{F_0}}.$$

**S.1 Validity of standard linear solid**

We use a test we developed in our previous publication (1) where we showed that it is possible to determine the viscoelastic model that better fits the data by plotting the data of *E′* against *E′′* *(The relationship should be linear for the SLS model, cf. Eq. 2 below)*. A scatter plot of $E'$ vs. $E^{0''}$ is shown in Figure S3; the plot confirms that the standard linear solid (SLS) is a good description for our system because the relation between $E'$ and $E''$ is nearly linear, as shown in (1). In fact, the Pearson correlation coefficients for the pixels of the different walls of the image shown in Fig. S3 are 0.96 for the transverse, 0.92, 0.91 and 0.94 for the longitudinal, and 0.89, 0.91 and 0.71 for the periclinal walls. The small deviations from a perfect correlation between $E'$ and $E''$ in the anticlinal walls show again that the standard linear solid model is a good approach but even the larger deviations in the periclinal walls should not be a problem, because even the generalised Maxwell model can always be described with a first-order approximation of the standard linear solid model.

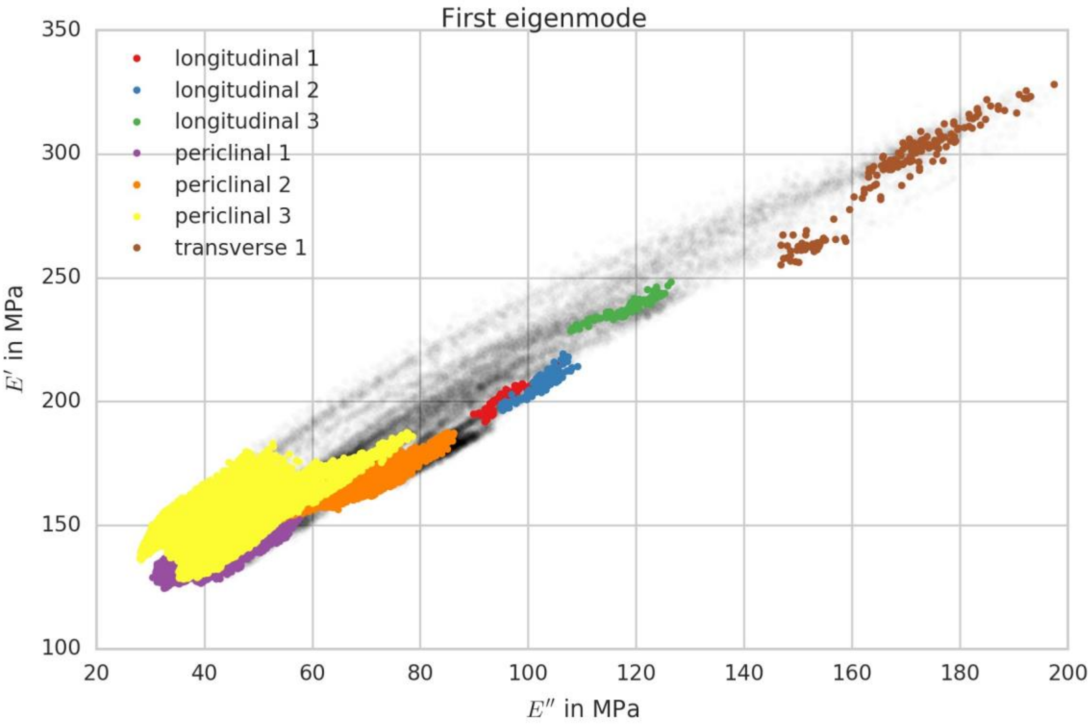


**Figure S3. Standard linear solid behaviour of measurements with the first eigenmode. The relation between $E'$ and $E''$ is an indicator of which viscoelastic model the material follows. The grey dots in the plot represent all data points from a dynamic AFM scan for $E'$ plotted against $E''$. The coloured dots represent the different walls. All dots that are not coloured, were neglected during the**

**analysis to obtain a sharp separation of the different walls, and they belong mostly to periclinal walls. If the average of the pixels, i.e. the majority of the pixels follows a linear function, then the standard linear solid is the appropriate viscoelastic model. For the first eigenmode, which corresponds to a frequency of $f_1 \approx$ 78kHz, as shown in this figure, the SLS is a good approach.**

### S.2 Theory to calculate the time response of the cell wall and numerical implementation

For the SLS model, we previously introduced in our analysis (1) the following relationships relating $E'$ and $E''$ :

$$E' = k_\infty + k_M \frac{\omega^2\tau^2}{1+\omega^2\tau^2}$$

$$E'' = k_M \frac{\omega\tau}{1+\omega^2\tau^2} \tag{S1}$$

In the SLS, $k_\infty$ denotes the elastic constant associated with the purely elastic spring, corresponding to the instantaneous (high-frequency) response of the material. The parameter $k_M$ is the modulus of the Maxwell element spring, which governs the amplitude of the viscoelastic contribution. The angular frequency $\omega$ is the frequency of the applied oscillatory deformation, and $\tau$ is the characteristic relaxation time of the Maxwell element, defined by the ratio of the dashpot viscosity to the spring modulus. Together, these parameters describe the frequency-dependent storage modulus $E'$ and loss modulus $E''$ of the material.

In order to calculate the time response $\tau$ of each pixel of the image, it is useful to treat individual pixels as distinct elements, each characterized by its own local mechanical properties, while exploiting the spatial correlations between neighbouring pixels to extract additional information about the system. Adjacent pixels are necessarily mechanically coupled, as the pixel size is comparable to the size of the indenter. Moreover, oscillatory indentation of the cell wall (CW) generates mechanical waves that propagate through the material, dynamically exciting neighbouring pixels as a result of the imposed oscillations. In addition, the plant CW can be reasonably approximated as a smooth, continuous material at the length scale of the measurement, without singularities. Under these assumptions, adjacent pixels can be described by the same viscoelastic model. Within the framework of a standard linear solid (SLS) model, the storage modulus E′ can be expressed as a function of the loss modulus E′′ by combining Eqs. (S1) above, which yields:

$$E' = k_\infty + E''\omega\tau \tag{S2}$$

which provides a direct local relationship between elastic and viscous responses under oscillatory loading. From this equation it can be readily shown that τ for each pixel can be obtained using the relation:

$$\tau = \frac{1}{\omega}\frac{\partial E'}{\partial E''} \tag{S3}$$

Using this approach, it is now possible also to calculate the rest of the viscoelastic quantities of the standard linear solid, namely $k_\infty$, $k_m$, and $\eta$, using the following equations:

$$k_\infty = E' - E''\omega\tau$$

$$k_M = E'' \frac{1+\omega^2\tau^2}{\omega\tau} \quad \text{(S4)}$$

$$\eta = \tau k_M$$

**Numerical implementation**

Equation S3 uses the spatial information to determine the relevant timescale of the material but the numerical derivative of $E'$ with respect to $E''$ cannot be implemented straight forwardly since it must be done independently with respect to the spatial dimensions. This assumes that the relevant timescale of the sample depends on the location, i.e. $\tau \equiv \tau(\boldsymbol{r})$, with $\boldsymbol{r} = (x,y)^T$ pointing to the location of the sample. Using this approach, we obtain with eqs. S1

$$\frac{\partial E'}{\partial \omega\tau} = \vec{\nabla} E' \frac{\partial \vec{r}}{\partial \omega\tau} = k_{\mathrm{M}} \frac{2\omega\tau}{(1+\omega^2\tau^2)^2} \quad \text{(S5a)}$$

$$\frac{\partial E''}{\partial \omega\tau} = \vec{\nabla} E'' \frac{\partial \vec{r}}{\partial \omega\tau} = k_{\mathrm{M}} \frac{1-\omega^2\tau^2}{(1+\omega^2\tau^2)^2} \quad \text{(S5b)}$$

,

with the Nabla operator $\nabla^{\sim} = \partial/\partial^{\sim} r$. Dividing now eq. S5a by eq. S5b, we obtain

$$\frac{\frac{\partial E'}{\partial \omega\tau}}{\frac{\partial E''}{\partial \omega\tau}} = \frac{2\omega\tau}{1-\omega^2\tau^2}. \quad \text{(S6)}$$

This can be brought into a quadratic equation and finally we obtain a solution

for $\tau$, which is given by

$$\tau = \frac{1}{\omega}\left(-D \pm \sqrt{D^2+4}\right), \quad \text{(S7)}$$

With $D = \left(\frac{\partial E'}{\partial \omega\tau} / \frac{\partial E''}{\partial \omega\tau}\right)^{-1}$.

The negative part of the solution can be neglected, because the relaxation time must be positive. Using this equation, it is now possible to calculate the time response for each pixel.

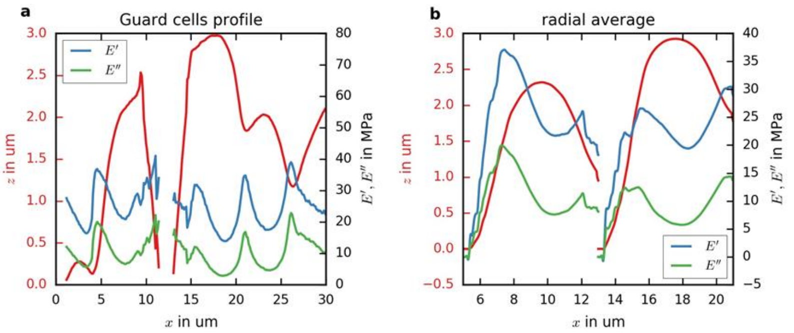


**Figure S4. Mechanical asymmetry in guard cells. Profiles of topography and dynamic moduli from the guard cells shown in Fig. 5. (a) Line profile along the dashed path in Fig. 5. (b) Averaged profiles for each guard cell, with the pore centred.**
